# Proteomic pipeline to identify age-related compositional and structural changes of the human dentin extracellular matrix

**DOI:** 10.1101/2020.05.27.116624

**Authors:** Mariana Reis, Fred Lee, Ana K. Bedran-Russo, Alexandra Naba

## Abstract

**Objective:** Devise a pipeline to investigate the protein composition of the human root dentin extracellular matrix (ECM) from single individuals of different age cohorts.

**Design:** Individual cervical root dentin of sound human molars from two age brackets, young (18-25 years old; n=3) and old (75-85 years old; n=3), were cut and pulverized. Protein extraction and fractionation were completed by sequential demineralization with EDTA buffer, chaotropic extraction with guanidine hydrochloride, and urea. The resulting protein extracts of differential solubility were digested into peptides and peptides were analyzed by mass spectrometry. Data generated for this study are available via ProteomeXchange, identifier PXD018320.

**Results:** We found that protein extracts of different solubilities present distinct biochemical compositions. We further define the matrisome of young (48 proteins) and old (50 proteins) human root dentin and report the identification of compositional and structural differences in ECM proteins from young and old teeth.

**Conclusion:** Our study provides a rigorous pipeline, from sample preparation to data analysis, to investigate the ECM composition – or matrisome – of the dentin. This pipeline has the potential to lead to the discovery of biomarkers of tooth aging and health.

## 1. Introduction

The world population is aging at a high rate. Studies predict that, by 2050, the number of adults over the age of 65 will increase by 16%, and the number of 80-year-olds and older individuals is estimated to triple (United Nations, Department of Economic and Social Affairs, 2019). These cohorts will sustain their natural dentition and experience higher prevalence of oral health conditions (Friedman et al., 2014). Particularly, root caries, a multifactorial disease on the root surface of the tooth compromises oral health and function in elderly individuals, with high prevalence in adults of 65 years old and above (Hayes et al., 2017; Zhang et al., 2020). This disease results in the progressive breakdown of the tissue as a consequence of mineral loss and degradation of the dentin extracellular matrix (ECM).

Mineralized tissues, like bone and teeth, are well-organized hierarchical structures consisting of an ECM that provides structural functionality for the deposition of an inorganic phase made of hydroxyapatite crystals (Alford et al., 2015; Marshall, 1993). With aging, the dentin tissue exhibits higher mineral content (Kinney et al., 2005), a gradual reduction of dentin tubule diameter and dental pulp size (Iezzi et al., 2019), and decreased mechanical properties (Arola & Reprogel, 2005; Ryou et al., 2015). However, there is a significant gap in our knowledge of the modifications in the composition and abundance of dentin ECM components with aging. Thus, having a cartography of the dentin ECM would be a step towards the discovery of potential biomarkers of the aging tooth. It would also be a step towards the development of strategies to maintain or regenerate a functional dentition.

Over the past decade, mass-spectrometry-based proteomic approaches have been used to study many aspects of oral biology including the profiling of the saliva and enamel pellicle (Lau et al., 2021; Rosa et al., 2012; Ventura et al., 2017) but also the composition of different parts of the tooth such as the enamel (Green et al., 2019; Hu et al., 2005; Jágr et al., 2019), the dentin (Horsophonphong et al., 2020; Jágr et al., 2012; Park et al., 2009; Widbiller et al., 2019), the pulp (Abbey et al., 2018; Eckhard et al., 2015; Jágr et al., 2016; Pääkkönen et al., 2005; Wei et al., 2008), and the cementum (Salmon et al., 2013). These studies employed diverse methodologies of sample preparation, protein extraction, identification, and quantification. For example, protein extraction was performed using different reagents, such as guanidine hydrochloride (Abbey et al., 2018; Eckhard et al., 2015; Jágr et al., 2012), radioimmunoprecipitation assay buffer (RIPA) (Park et al., 2009), ethylenediamine tetraacetic acid (EDTA), among others (Chun et al., 2011; Horsophonphong et al., 2020; Jágr et al., 2012; Sharma et al., 2020; Widbiller et al., 2019; Wojtas et al., 2020). The analysis of proteomic datasets has been explored using various bioinformatic tools that are critical to the interpretation of the MS/MS spectra and protein identification (Calderón-González et al., 2016; Cho, 2007).

However, while these studies identified a variety of proteins in human teeth, in many instances the level of confidence with which ECM proteins were identified lacked the rigor expected from such studies, and they were not designed to investigate specifically ECM proteins that present unique biochemical features. Moreover, among the fifteen studies assessed, and that focused on tooth proteomics, only five used human dentin as their substrate (Horsophonphong et al., 2020; Jágr et al., 2012, 2019; Park et al., 2009; Widbiller et al., 2019), and just two studies targeted the dentin from the root of the tooth (Jágr et al., 2012; Park et al., 2009). Interestingly, only one study investigated separately the protein composition of young individuals (n=3) (Park et al., 2009), while the majority of the studies pooled their samples, regardless of age, gender, or area of the tooth. Furthermore, the lack of use of a unified annotation of ECM components prevented a comprehensive definition of the dentin ECM proteins and comparisons across different cohorts and studies.

The large size and high insolubility of ECM proteins have made their biochemical analysis notoriously difficult (Taha & Naba, 2019). In the past years, we and others have developed experimental proteomic pipelines tailored to enrich, solubilize, and digest ECM proteins (Naba et al., 2012; Randles & Lennon, 2015; Schiller et al., 2015; Taha & Naba, 2019). We have also used bioinformatics to define the “matrisome”, the collection of genes encoding ECM and ECM-associated proteins, and have shown that the matrisome list can be used to comprehensively annotate high-throughput data for ECM components (Gebauer & Naba, 2020; Naba et al., 2016; Naba & Ricard-Blum, 2020). With these tools in place, proteomics has now become a method of choice to study the changes in the ECM associated with diseases such as cancers (Socovich & Naba, 2019), fibrosis (Schiller et al., 2019) or during aging (Angelidis et al., 2019; Grilo et al., 2020).

Here, we assessed the feasibility of using proteomics to characterize the ECM composition of the root dentin from single individuals. To do so, we devised an experimental and stringent analytical pipeline to compare the ECM proteins found in the soluble and insoluble components of the human cervical root dentin. Using this pipeline, we first assessed the extent of inter-individual variability in cohorts of same age and gender. We then compared the ECM protein composition of root dentin from young and older adult individuals. Last, we showed that we could gain structural insight from the proteomic data. Altogether, our study provides a pipeline to investigate the composition of the dentin ECM that has the potential to lead to the discovery of biomarkers of the aging tooth.

## 2. Material and methods

### 2.1. Sample selection and preparation

Three sound molars were selected from two age brackets, young (18-25 years old) and older (75-85 years old) men, according to protocol 2018-0346 approved by the Institutional Review Board Committee of the University of Illinois at Chicago (**Supplementary Table 1A**). Teeth were immediately stored after extraction at −20°C. The cervical part of the root (**Fig. 1A**) was obtained by sectioning the tooth with an IsoMet™ high precision cutting machine (Buehler Ltd, Lake Bluff, IL, USA). All soft tissues, periodontal ligament, cementum, and pulp tissue were carefully removed with curettes and endodontic files.

**Figure 1.**
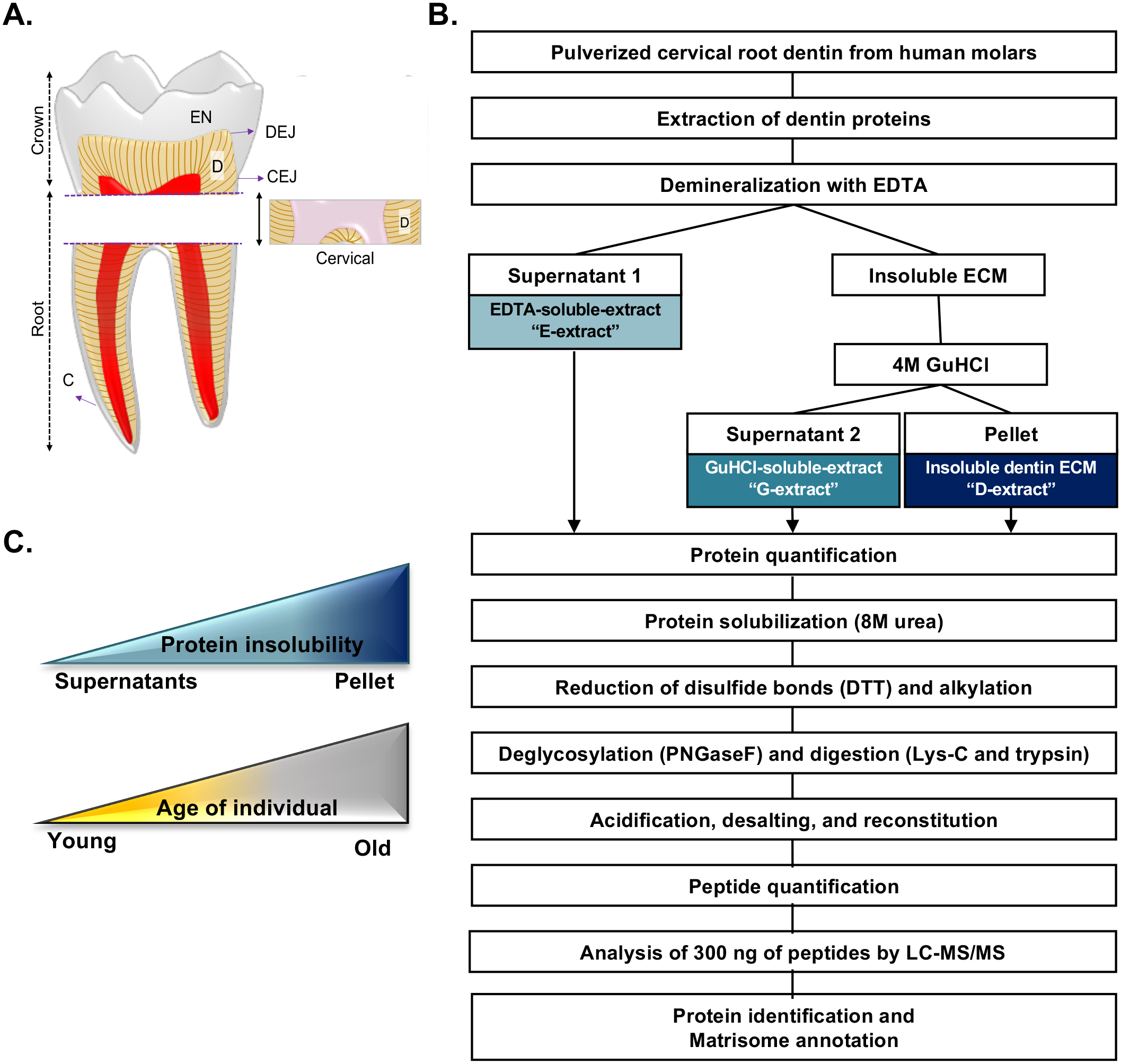
Protein fractionation pipeline for the cervical root dentin. Schematic of a sound human molar (EN: enamel, D: dentin, C: cementum, DEJ: dentin-enamel junction, CEJ: cement-enamel junction, and P: pulp) and the type of dentin (root) and area (cervical) used for protein extraction (**A**). Experimental pipeline describing sample preparation, protein extraction, digestion, and mass spectrometry analysis (**B**). Color codes used throughout the figures to identify the different protein extracts and age of the individuals (**C**).

### 2.2. Dentin protein extraction

The extraction of dentin proteins was adapted from an existing protocol (Martin-De Las Heras et al., 2000; Park et al., 2009), with modifications (**Fig. 1B**). In brief, the cervical dentin root sections (250-350 mg per tooth/sample) were pulverized, and the powder demineralized with a 0.5M EDTA solution at pH 8.0 containing protease inhibitors (Pierce Protease Inhibitor Tablets, EDTA-free, Thermo Fisher Scientific, Rockford, IL, USA) for 12 days under agitation at 4°C. The EDTA solution was changed every 3 days. Solubilized proteins were obtained by centrifugation at 5000 x g for 10 min at 4°C. The resulting supernatant containing EDTA-soluble proteins will be subsequently referred to as the “E-extract”. Protein extraction from the remaining EDTA-insoluble pellet was performed with a 4M guanidine hydrochloride (Gu-HCl, Sigma-Aldrich St. Louis, MO, USA) solution at pH 7.4 containing protease inhibitors, for 3 days under agitation at 4°C. Solubilized proteins were obtained by centrifugation at 5000 x g for 10 min at 4°C. The resulting supernatant containing Gu-HCl-soluble proteins will be subsequently referred to as the “G-extract”. Proteins from the remaining Gu-HCl-insoluble pellet will be further termed the dentin extract or “D-extract”. The E- and G-extracts were dialyzed against ultrapure water using M_r_ 6,000 cut-off Spectra/Por dialyses tubing (Spectrum Laboratories, Rancho Dominguez, CA, USA) at 4°C for 6 h. The ultrapure water was exchanged twice. The total concentration of proteins was measured with the Pierce Rapid Gold BCA Protein assay kit (Thermo Fisher Scientific, Rockford, IL, USA). The E-, G-, and D-extracts were lyophilized (FreeZone Freeze Dryer, Labconco, Kansas City, MO, USA) and stored at −20°C for further analyses.

### 2.3. Protein digestion

The lyophilized samples (E-extracts, G-extracts and D-extracts, 5-10 mg dry weight) were subsequently solubilized and digested into peptides following an established and previously described protocol (**Fig. 1B**) (Naba et al., 2012). Briefly, the proteins were solubilized in 8M urea, and protein disulfide bonds were reduced using 10mM dithiothreitol and alkylated with 25mM iodoacetamide (Thermo Fisher Scientific, Rockford, IL, USA). Proteins were then deglycosylated with PNGaseF (New England Biolabs, Ipswich, MA, USA) and digested with Lys-C (Thermo Fisher Scientific, Rockford, IL, USA) for 2 hours at 37°C, and trypsin (Thermo Fisher Scientific, Rockford, IL, USA), overnight at 37°C. The samples were acidified with 50% trifluoroacetic acid (TFA, Thermo Fisher Scientific, Rockford, IL, USA) until the pH of the samples was lesser than 2. Acidified samples were desalted in C18 desalting columns (Pierce Peptide Desalting Spin Columns, Thermo Fisher Scientific, Rockford, IL, USA) and reconstituted in 95% HPLC grade water, 5% acetonitrile and 0.1% formic acid. Peptide concentration was measured with a colorimetric assay (Pierce Quantitative Colorimetric Peptide Assay kit, Thermo Fisher Scientific, Rockford, IL, USA).

### 2.4 Analysis of digested dentin proteins by LC-MS/MS

Approximately 300 ng of desalted peptides were analyzed at the University of Illinois Mass Spectrometry core facility on a Thermo Fisher Orbitrap Velos Pro coupled with Agilent NanoLC system (Agilent, Santa Clara, CA). The LC columns (15 cm × 75 μm ID, Zorbax 300SB-C18) were purchased from Agilent. Samples were analyzed with a 120-min linear gradient (0–35% acetonitrile with 0.1% formic acid) and data were acquired in a data-dependent manner, in which MS/MS fragmentation was performed on the top 10 intense peaks of every full MS scan. Full MS scans were acquired in the Orbitrap mass analyzer over m/z 350–1800 range with resolution 30,000 (m/z 400). The target value was 1.00E+06. The ten most intense peaks with charge state ≥ 2 were fragmented in the HCD collision cell with normalized collision energy of 35%, these peaks were then excluded for 30 s after 2 counts within a mass window of 10 ppm. Tandem mass spectrum was acquired in the Orbitrap mass analyzer with a resolution of 7,500. The target value was 5.00E+04. The ion selection threshold was 5,000 counts, and the maximum allowed ion accumulation times were 500 ms for full scans and 250 ms for HCD.

All MS/MS samples were analyzed using Mascot (Matrix Science, London, UK; version 2.6.2). Mascot was set up to search the Uniprot-human_20190917 database (20,430 entries; individuals 1 and 6) Uniprot-human_20191015 database (20,365 entries; individuals 2, 3, 4, 5) assuming stricttrypsin. Mascot was searched with a fragment ion mass tolerance of 0.30 Da and a parent ion tolerance of 15 PPM. Carbamidomethyl of cysteine was specified in Mascot as a fixed modification. Gln->pyro-Glu of the N-terminus, deamidated of asparagine and glutamine and oxidation (hydroxylation) of methionine, proline, and lysine were specified in Mascot as variable modifications.

Scaffold (version Scaffold_5.0.1, Proteome Software Inc., Portland, OR) was used to validate MS/MS based peptide and protein identifications. The following stringent thresholds were applied: peptide identifications were accepted if they could be established with a false-discovery rate (FDR) of less than 1.0% by the Peptide Prophet algorithm (Keller et al., 2002) with Scaffold delta-mass correction. Protein identifications were accepted if they could be established at greater than 95.0% probability and contained at least 2 identified peptide. Protein probabilities were assigned by the Protein Prophet algorithm (Nesvizhskii et al., 2003). Proteins that contained similar peptides and could not be differentiated based on MS/MS analysis alone were grouped to satisfy the principles of parsimony. Proteins sharing significant peptide evidence were grouped into clusters.

Mass spectrometry output was further annotated to identify ECM and non-ECM components using the human matrisome list (Naba et al., 2012, 2016). In brief, matrisome components are classified as core-matrisome or matrisome-associated components, and further categorized into groups based on structural and/or functional features: ECM glycoproteins, collagens or proteoglycans for core matrisome components; and ECM-affiliated proteins, ECM regulators, or secreted factors for matrisome-associated components (Hynes & Naba, 2012; Naba et al., 2012).

Raw mass spectrometry data have been deposited to the ProteomeXchange Consortium (Deutsch et al., 2020) via the PRIDE partner repository (Perez-Riverol et al., 2019) with the dataset identifier PXD018320 and 10.6019/PXD018320. *The raw data will be made publicly available upon acceptance of the manuscript*.

## 3. Results

### 3.1. Feasibility of the analysis of samples from single individuals

To assess the feasibility of using proteomics to characterize the ECM composition of root dentin from each individual, we used the cervical area of the root of sound molars from young individuals (18-25 years old) and fractionated it to generate protein samples of different solubility (**Fig. 1B**). The sample preparation protocol yielded, for each sample, three distinct fractions, an EDTA-soluble-extract “E-extract”, a Gu-HCl-soluble-extract “G-extract”, and a remaining pool of insoluble dentin proteins, enriched for ECM proteins, further termed the “D-extract” (**Fig. 1B**). A total of 5-10 mg (in dry weight) of each protein extract was solubilized and digested into peptides as previously described (Naba et al., 2012). After digestion and desalting, the amount of peptide obtained for each type of extracts ranged from of 10-140 µg for the E-extracts, 13-189 µg for the G-extracts and 155-343 µg for the D-extracts. We observed that the least amount of peptides were obtained from the most soluble E-extract while the most insoluble protein fraction, D-extract, contained the most peptides. Nevertheless, sufficient amount of peptides were obtained from each extract for mass spectrometric analysis. This demonstrates the feasibility of our pipeline, where a sample from a single tooth and from a single individual provides enough material for the downstream proteomic analysis.

While many of the dentin proteomic studies focused on the analysis of the most soluble protein fractions, the goal of this study was to evaluate both soluble and insoluble components, and thus assess the inter-individual variability in the protein composition and solubility of the human cervical root dentin within cohorts of same age and gender. To streamline the reporting of our results, we further combined the identified protein fractions of higher solubilities, the E- and G-extracts, into one larger group subsequently termed “supernatant” and compared these lists to the ECM proteins identified in the insoluble dentin pellet (*i.e.,* resistant to EDTA and Gu-HCl solubilization), the “D-extract”.

### 3.2. Protein composition of extracts of different solubilities

By considering ECM proteins detected with at least two peptides in at least one sample, a total of 99 proteins were detected in the cervical area of root dentin of young individuals (**Supplementary Table 1B**), 48 of which were matrisome proteins (**Supplementary Tables 1C and 1D**). Initially, we compared the matrisome proteins of supernatants and D-extracts from young dentin samples (n = 3). We found 38, 42 and 39 ECM proteins from individuals 1, 2, and 3, respectively (**Fig. 2A and Supplementary Tables 1D-G**). Collectively, supernatants (33-38 proteins) presented a higher number of identified ECM proteins than D-extracts (24-27 proteins) The number of ECM proteins detected in both supernatants and D-extracts was similar among the three individuals, an average of 21 proteins (**Supplementary Tables 1D-G**). We also observed that proteins fractions of higher solubility, *i.e.,* the supernatants, contained a higher number of uniquely identified ECM proteins, 12 to 17 (36-45%), in comparison to the D-extracts (2 to 6 proteins or 8-22% of the proteins detected) (**Fig. 2A**). The analysis of the insoluble D-extracts revealed that 50% of the identified ECM proteins are collagens and that half as many glycoproteins, proteoglycans, and secreted proteins were detected compared to the supernatants (**Supplementary Tables 1D-G**).

**Figure 2.**
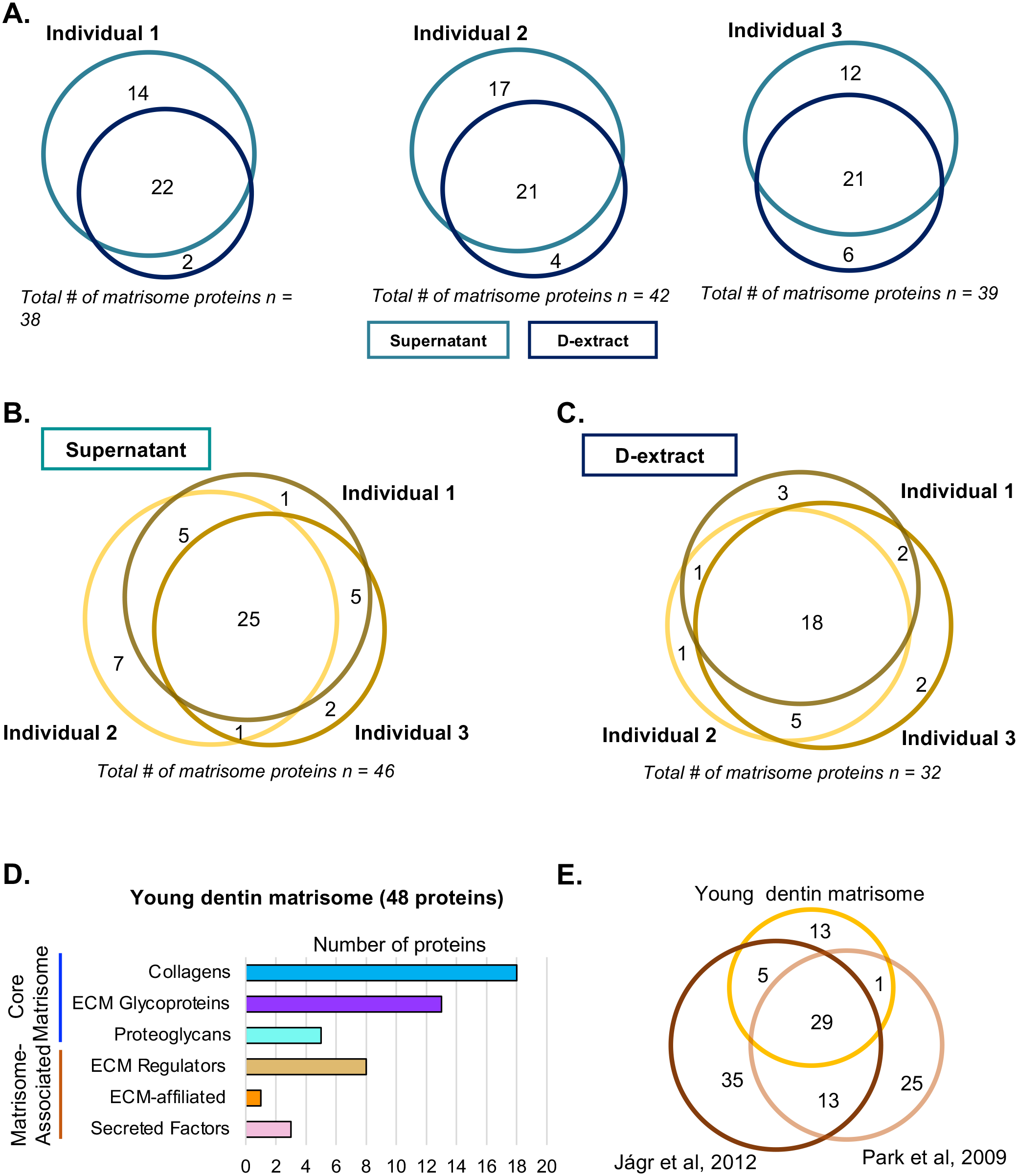
Profiling the young dentin matrisome. Venn diagrams present the number of matrisome proteins detected in supernatants (combined E- and G-extracts) and D-extracts of young individuals (**A**). Venn diagrams present the inter-individual comparisons of young ECM proteins detected in fractions of different solubility: supernatants (**B**) and D-extracts (**C**). Bar chart shows the distribution of proteins detected in young dentin across the different matrisome categories. Proteins included in the analysis were detected with at least two peptides. The young dentin matrisome is defined as the ensemble of proteins detected in at least one of the fractions and individual (**D**). Venn diagram illustrates the number of shared and distinct proteins between the lists defined as the young dentin matrisome in this study and the re-annotated proteomes of human dentin published by Park et al., 2009 and Jágr et al., 2012 (**E *and Supplementary Table 2***).

### 3.3. Assessment of inter-individual variability

We next sought to assess the extent of inter-individual variability of dentin proteins between three distinct young individuals. We collated the proteins identified in either supernatants or D-extracts for each individual and compared these lists (**Fig. 2B-C and Supplementary Tables 1D-G**). The union of three lists revealed 46 and 32 ECM proteins were identified respectively in the supernatants and D-extracts of young individuals (**Fig. 2B-C and Supplementary Table 1H**). We identified a set of 25 ECM proteins in supernatants that intersect across all individuals (**Fig. 2B**), and 18 common ECM proteins in D-extracts (**Fig. 2C**). The number of uniquely identified ECM proteins found in each young individual varied according to the fraction, where a higher number of proteins was detected in supernatants (10), compared to D-extracts (6), indicating a greater variability in samples of higher solubility.

### 3.4. Defining the young dentin matrisome

To further characterize the ECM composition of the young cervical dentin, we defined the dentin “matrisome” as the ensemble of proteins detected in at least one individual and in either supernatants or D-extracts. Using this definition, the dentin matrisome of young adult individuals comprises of 48 ECM proteins (**Fig. 2D, Table 1, Supplementary Table 1H**). The dentin ECM composition of the young cohort consisted of predominantly 18 collagens (38%) and 13 glycoproteins (27%), along with 5 proteoglycans (10%), where, together, they constitute the core matrisome. Matrisome-associated proteins were also present, however in a smaller quantity, represented by 1 ECM-affiliated protein (2%), 8 ECM regulators (17%), and 3 secreted factors (6%) (**Fig. 2D**).

**Table 1.**
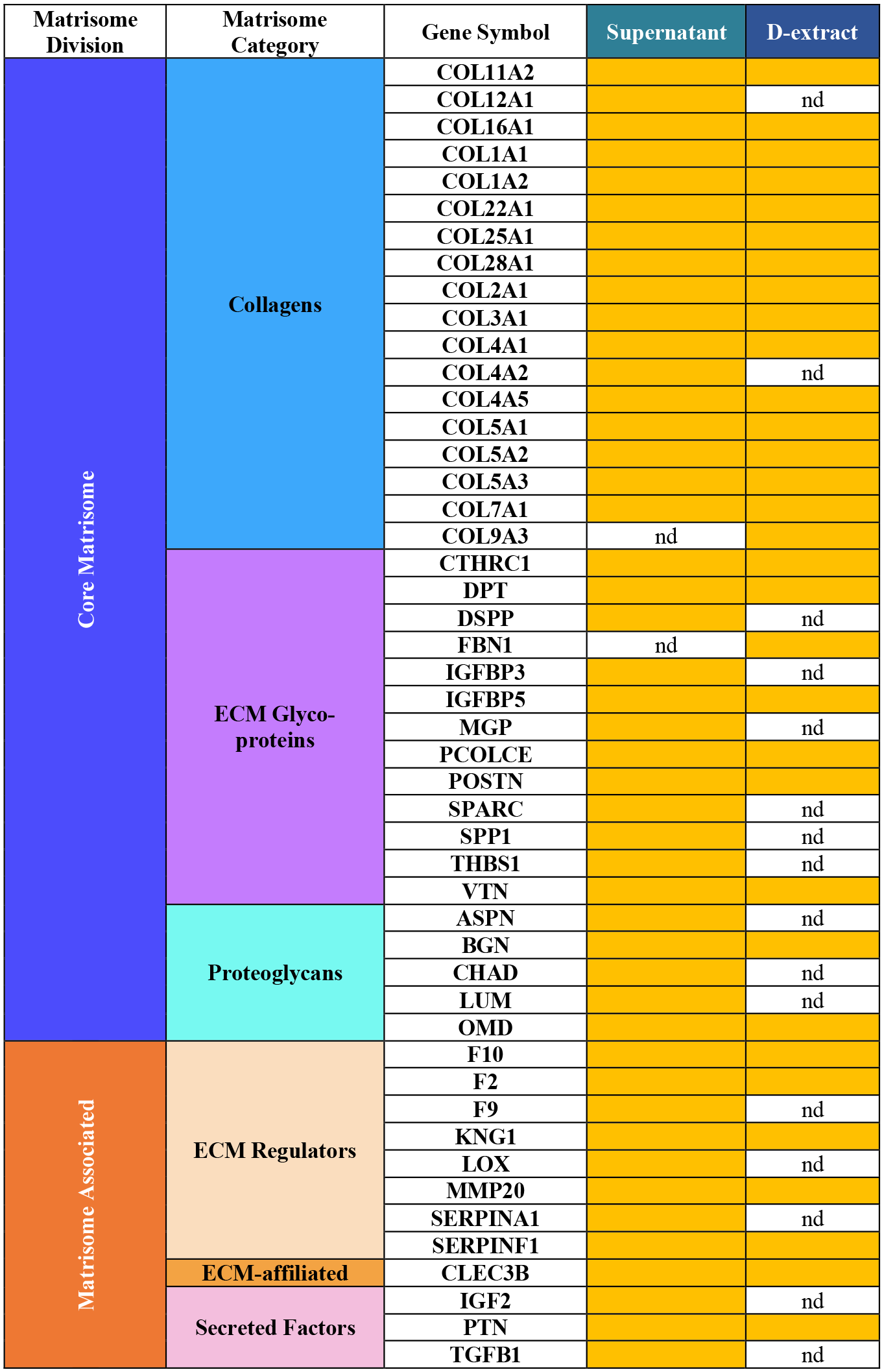
List of ECM proteins of the young dentin matrisome. The proteins included in the list were detected with at least 2 peptides and in one or more individuals. Data are sorted by fractions and matrisome categories (nd: not detected).

The main proteins that have previously been shown to compose the dentin ECM were detected. Collagens, as expected, were the most abundant protein in the young dentin matrisome. The two chains forming the fibrillar type I collagen (COL1A1 and COL1A2) were predominant and the most abundant protein detected in our sample (**Supplementary Table 1D, columns O – W** representing normalized total spectral count, a proxy for protein abundance), however, we also report the identification of a number of collagens not previously associated with the dentin tissue, including collagens type IV-V, VII, IX, XVI, XXII, XXV, and XXVIII (**Table 1 and Supplementary Table 1H**).

We identified glycoproteins such as dentin sialophosphoprotein (DSPP), osteopontin (SPP1), procollagen C-endopeptidase enhancer (PCOLCE), secreted protein acidic and cysteine rich (SPARC), matrix Gla protein (MGP) and vitronectin (VTN) that are known dentin proteins with important roles in biomineralization, enzymatic cleavage of type I procollagen and cell adhesion (Kawasaki et al., 2004; Linde, 1989; Ravindran & George, 2014; Ritchie, 2018). We also report the detection of other glycoproteins never found before in the root dentin ECM, such as collagen triple helix repeat-containing protein 1 (CTHRC1) and fibrillin 1 (FBN1), the later detected in both supernatant and D-extracts, and the last exclusively in D-extracts.

Small leucine-rich proteoglycan (SLRP) family were the major proteoglycans (PGs) found. They include biglycan (BGN) and asporin (ASPN) (Class I), lumican (LUM) and osteomodulin (OMD) (Class II), and chondroadherin precursor (CHAD) (Class IV), which are known to participate in collagen fibril assembly, tissue growth and regulation of interfibrillar spacings (Iozzo & Schaefer, 2015).

ECM-associated proteins were identified including the coagulation factors (F2, F9 and F10), and a matrix metalloproteinase protein (MMP20) that are involved in the breakdown of extracellular matrix in normal physiological processes, tissue remodeling/degradation, and disease processes (Loffek et al., 2011). Insulin like growth factor 2 (IGF2), a member of the insulin family of polypeptide growth factors and involved in development and growth of tissues (Baxter, 2000) was identified for the first time in the root dentin.

We further aimed to compare the list of ECM proteins identified here with those reported in previous investigations that explored the proteome of human root dentin (Jágr et al., 2012; Park et al., 2009). To compare our data with the earlier studies, we retrieved and reannotated them with the *in silico*-predicted complete matrisome list (Naba et al., 2016). The data are presented in **Supplementary Table 2A**. The assessment revealed 29 ECM proteins found in all three studies and another set of 19 ECM proteins that were detected in two out of the three studies (**Fig. 2E and Supplementary Table 2B**). The presence of proteins such as dermatopontin (DPT), dentin sialophosphoprotein (DSPP), biglycan (BGN), and matrix metallopeptidase-20 (MMP20) further confirms the established association of these proteins to the dentin. However, a greater difference is revealed between the three matrisomes: while 35 and 25 proteins were unique to the two studies (Jágr et al., 2012; Park et al., 2009), respectively, our study identified an additional 13 ECM proteins not detected before in the human root dentin. These 13 structural ECM proteins consisted of 10 collagens, 2 glycoproteins and 1 secreted factor similarly present in young supernatants and D-extracts (**Fig. 2E and Supplementary Table 2**). This meta-analysis demonstrates the strength of our ECM-focused proteomic and analytical pipeline in the cervical area of root dentin and reveals an additional list of collagens never identified in this type of sample.

Since our approach allowed for the identification of proteins from individual dentin samples of different solubilities, we further compared the samples of similar solubilities from the 2012 study by Jágr et al. to our respective E-, G-, and D-extracts. (**Fig. S1 and Supplementary Tables 2C-E**). A set of 22 ECM proteins were identified in E-extracts of both studies (**Fig. S1A**). G-extracts and D-extracts revealed 22 and 11 ECM common proteins, respectively (**Fig. S1B-C**). While the comparison of G-extracts revealed a similar number of proteins that were uniquely identified in each study (22 – young matrisome and 33 – Jágr et al., 2012), the D-extract of the young matrisome, contained a significantly greater number of ECM proteins identified (21) when compared to Jágr et al (1), (**Fig. S1C**). The compositional differences noticed in the D-extracts are remarkable and indicate that advances in the enrichment of the ECM and protein solubilization might have had a significant impact on protein identification.

Of note, the dentin tissue is part of the dentin-pulp complex, where the odontoblast layer forms the outermost layer of the pulp and is adjacent to the predentin, an initial stage of new dentin tissue. Odontoblast processes extend into the dentinal tubules (Pashley, 1996). As such, proteins from this tissue could overlap with the dentin-pulp complex tissue, and even possibly contaminate the dentin tissue. We thus sought to compare the young dentin matrisome with data from prior studies on the proteome of the pulp tissue and/or odontoblast layer (Abbey et al., 2018; Pääkkönen et al., 2005; Wei et al., 2008). The comparison with Pääkkönen study on pulp showed only one protein, SERPINA1, was also found in the young dentin matrisome defined here, while we found no overlap between our study and the Wei et al., 2008 study. A total of 153 matrisome proteins were identified as part of the pulp and 102 matrisome proteins as part of the odontoblast layer, 84 of these being found in both compartments (Abbey et al., 2018). Our comparison revealed that 27 of the 153 proteins found in the pulp were also part of the young dentin matrisome and 24 of the 102 proteins of the odontoblast layer overlapped with the young dentin matrisome (**Fig. S1D and Supplementary Table 2F**). Importantly, 18 proteins of the young dentin matrisome were neither identified in the pulp nor the odontoblast layer. These comparisons demonstrate that there are distinct ECM signatures that define the different layers of the tooth even though a substantial association is seen in the tissues of the dentin-pulp complex.

### 3.5. Defining the old dentin matrisome

With our pipeline in place, we next compared the ECM composition of the root dentin of young individuals (*see above*) to that of older individuals. To do so, we analyzed samples from 3 older adults (75-85 years old). A total of 101 proteins in the cervical area of root dentin of old individuals was identified (**Supplementary Table 1B**), where 50 were matrisome proteins (**Supplementary Tables 1C and 1I**). The total number of ECM proteins identified from individuals 4-6 was 43, 40 and 45, respectively (**Fig. 3A and Supplementary Tables I-L**). The number of shared ECM proteins was comparable between individuals, where an average of 16 proteins identified in supernatants and D-extracts. Conversely, the number of ECM proteins found in either fraction varied according to the fraction and individual. The supernatants of two out of three old individuals showed a higher number of unique ECM proteins, 24 - 27 (55%), in comparison to 3 unique ECM proteins (7%) detected in the D-extracts (**Fig. 3A**). Individual 5 showed in opposing trend, however with a comparable number of total and overlapping ECM proteins between fractions.

**Figure 3.**
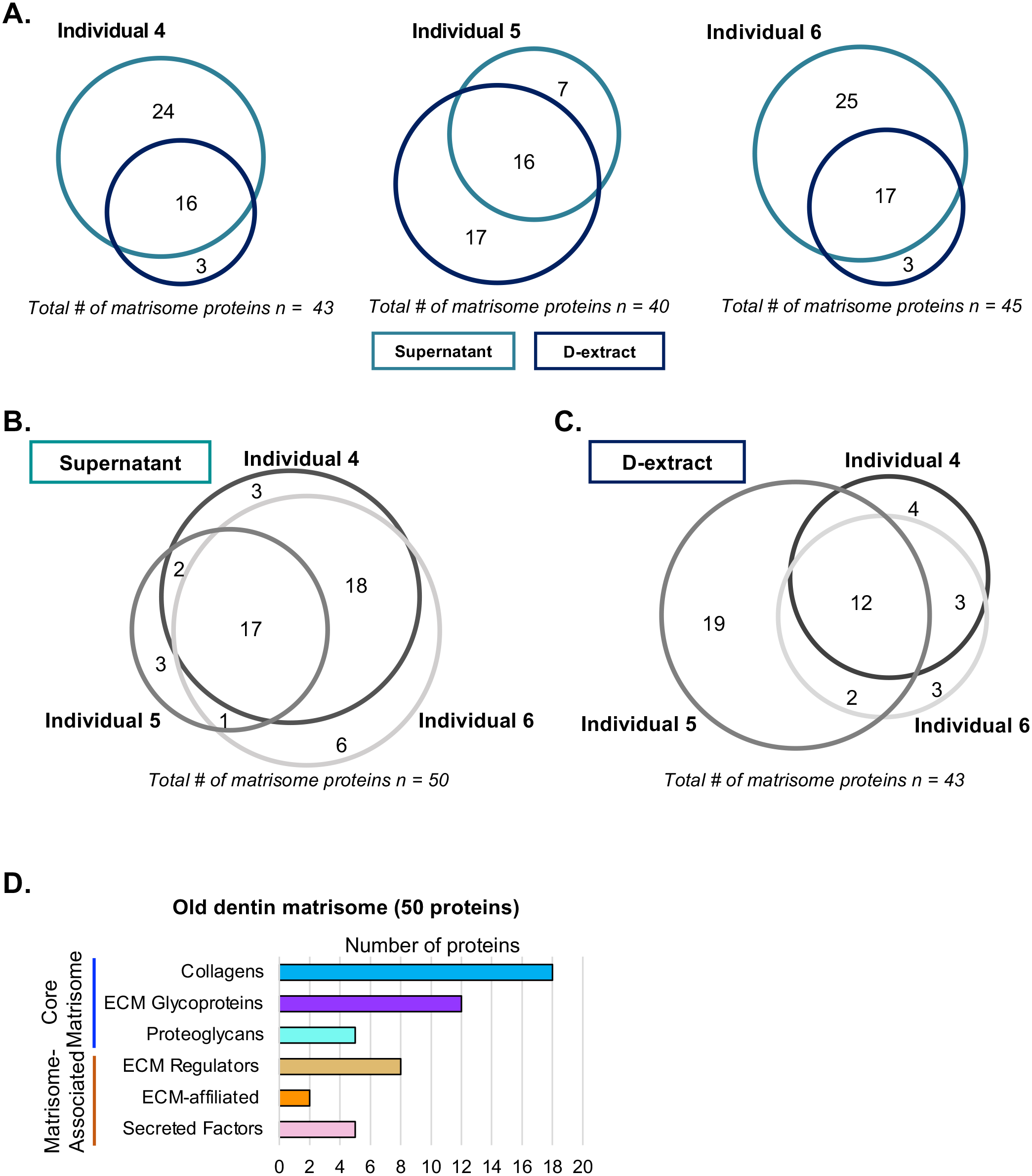
Profiling the old dentin matrisome. Venn diagrams present the number of matrisome proteins detected in supernatants (combined E- and G-extracts) and D-extracts in old individuals (**A**). Venn diagrams present the inter-individual comparisons of old ECM proteins detected in fractions of different solubility: supernatants (**B**) and D-extracts (**C**). Bar chart shows the distribution of proteins detected in old dentin across the different matrisome categories. Proteins included in the analysis were detected with at least two peptides. The old dentin matrisome is defined as the ensemble of proteins detected in at least one of the fractions and individual (**D**).

The inter-individual variability across older samples was evaluated and we identified ECM proteins in either supernatants or D-extracts for each individual and compared these lists (**Fig. 3B**). Our analysis identified 50 proteins in the supernatants (**Fig. 3B**) and 43 in the D-extracts (**Fig. 3C**). A set of 17 ECM proteins was identified in supernatants that overlap between all individuals, and 12 ECM common proteins in D-extracts. The number of ECM proteins uniquely identified in older individuals varied given the fraction, where a higher number of ECM proteins was detected in D-extracts (26), compared to those in the supernatants (12), revealing an increased variability in samples of higher insolubility.

Last, we defined the old dentin matrisome as the ensemble of proteins detected in at least one individual and in either supernatants or D-extracts that comprises of a total of 50 ECM proteins (**Table 2 and Supplementary Table M**). The old matrisome is thus constituted of a majority of core matrisome proteins, 18 collagens (36%), 12 ECM glycoproteins (24%) and 5 proteoglycans (10%). Fibrillar type I collagen (COL1A1 and COL1A2) was the most abundant protein detected in our sample (**Supplementary Table 1I, columns O – W** representing normalized total spectral count, a proxy for protein abundance). The matrisome-associated proteins were less evident and comprised 2 ECM-affiliated proteins (4%), 8 regulators (16%), and 5 secreted factors (10%) (**Fig. 3D and Table 2**).

**Table 2.**
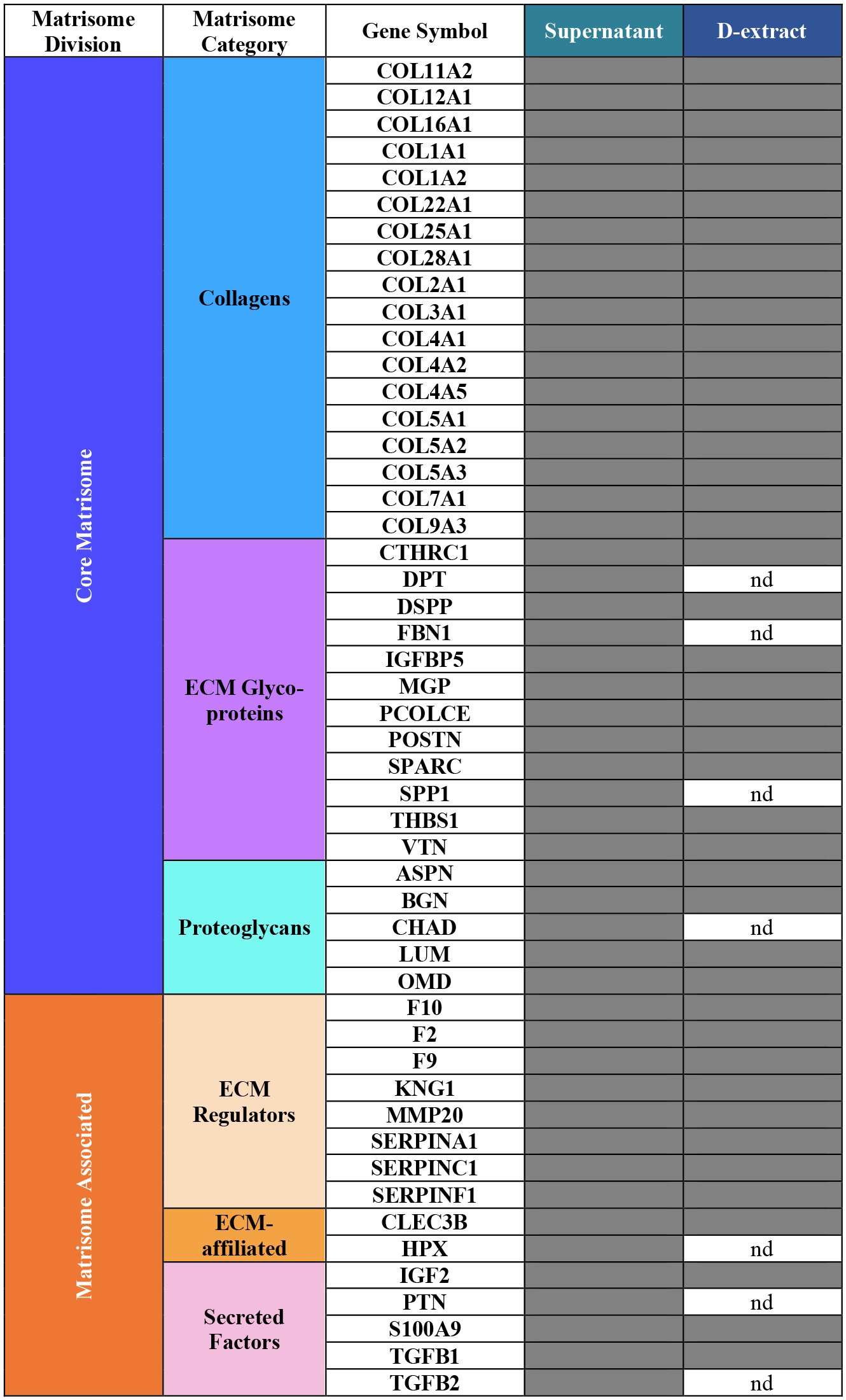
List of ECM proteins of the old dentin matrisome. The proteins included in the list were detected with at least 2 peptides and in one or more individuals. Data are sorted by extracts and matrisome categories (nd: not detected).

### 3.6. Comparing the root dentin ECM of young and older individuals

Having defined the “old root dentin matrisome”, our dataset now allows us to address the following questions: 1) can we identify changes between the ECM composition of young and old root dentin; 2) is there a difference in the solubility of the ECM proteins of young and older root dentin; and 3) can proteomic data, in particular analysis of sequence coverage, be used to infer difference in ECM protein digestibility, which could be used as a proxy for evaluating structural accessibility and protein folding.

#### 3.6.1. Comparing the ECM protein composition of young and old dentin samples

The comparison of the two age groups revealed that a large portion of ECM proteins (46) are identified in young and old samples (**Fig. 4A-B**). The major components found in both groups were 17 collagens (39%) and 11 ECM glycoproteins (26%). A smaller number of 5 proteoglycans (11% of the matrisome proteins) and ECM-affiliated proteins (1) and secreted factors (3) were detected. Overall, core matrisome proteins (33) were predominant (76%) in the ECM of both age groups, when compared to matrisome-associated proteins, 16 (15%). A set of 2 and 4 ECM proteins were uniquely found in young and old cervical root dentin, respectively. **Fig. 4A** shows the ECM proteins uniquely identified in young and old dentin matrisomes. Insulin-like growth factor-binding protein 3 (IGFBP3) and lysyl oxidase (LOX), were exclusively found in the young matrisome, while hemopexin (HPX), antithrombin-III (SERPINC1), protein S100-A9 (S100A9) and transforming growth factor beta-2 proprotein (TGFB2) were uniquely found in the old matrisome. The comparison between young and old matrisomes ultimately reveals that, while sharing a large number of proteins (46), each matrisome presents certain unique components that may relate to the particular functions of the dentin tissue. We can anticipate that expanding the sample size, may lead to the identification of additional proteins, whose abundance change with age. Beyond identification, our proteomic approach can reveal semi-quantitative differences in protein abundance (see columns reporting normalized total spectral counts in **Supplementary Tables 1B, C, D and I**). While our analysis did not reveal significant variation in abundance with aging, quantitative proteomic methods, such as those relying on peptide labeling, could provide further insights.

**Figure 4.**
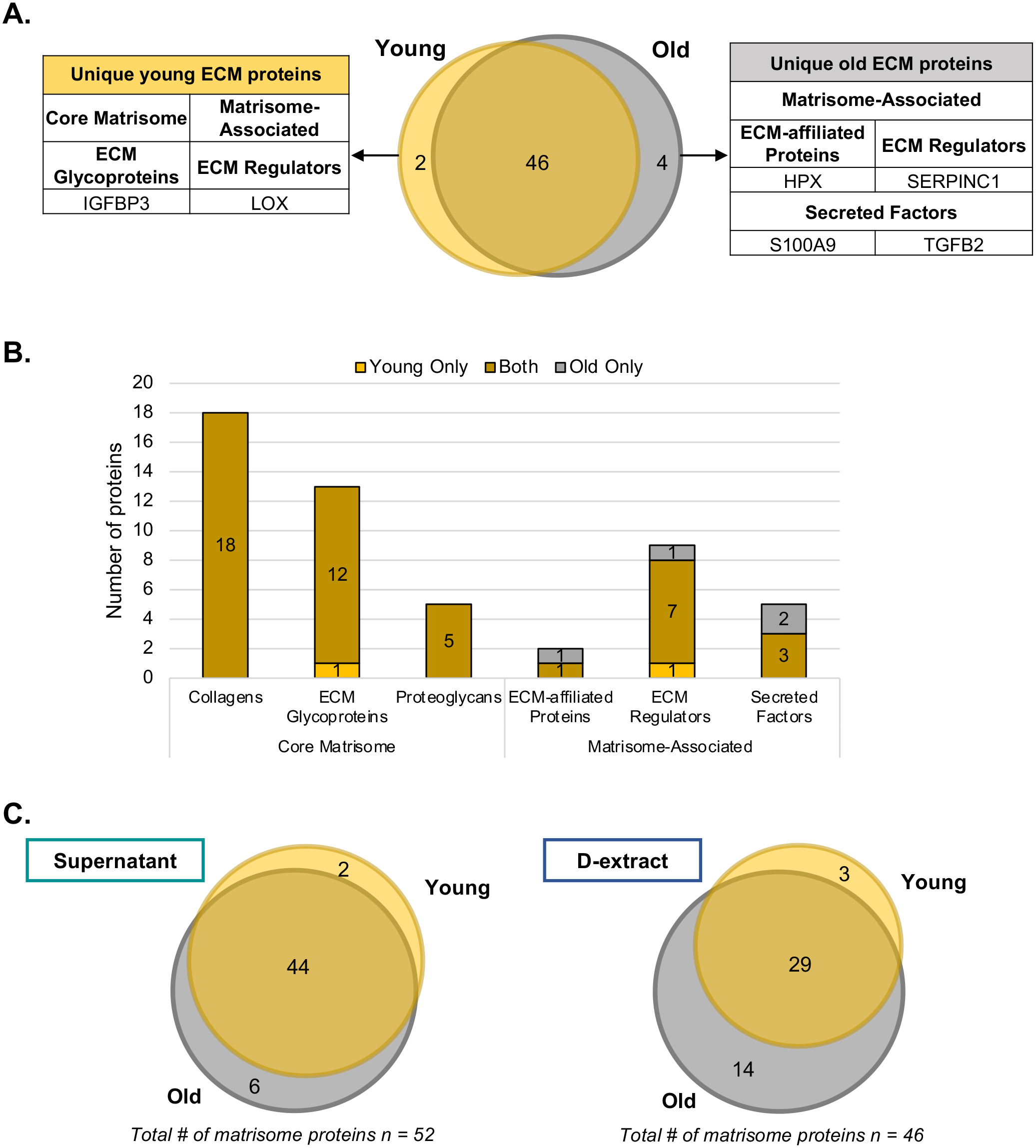
Comparison of the young and old dentin matrisomes. Venn diagram illustrates the number of matrisome proteins detected in either or both age groups. Lists of ECM proteins uniquely identified in young (left panel) and old (right panel) individuals (**A**). Bar chart shows the distribution of proteins detected in young (yellow) and old (grey) dentin matrisomes across the different matrisome categories (**B**). Venn diagrams present the comparisons of young and old ECM proteins detected in fractions of different solubility (supernatants and D-extracts) (**C**).

#### 3.6.2. Comparing the ECM protein solubility of young and old dentin samples

Our results also show that a higher number of proteins are identified in the more soluble protein fractions (supernatants), regardless of the age group, when compared to the more insoluble fractions (D-extracts). Additionally, we found that half or more ECM proteins are common to both type of samples. A large fraction of matrisome proteins in supernatants (44) are detected in both young and old cohorts (**Fig. 4C**). The main protein category identified was collagen, represented by 17 different collagens of the total proteins (38.6%). ECM glycoproteins accounted for 11 of the shared proteins (25%), where insulin like growth factor binding protein 3 (IGFBP3) and fibrillin 1 (FBN1) were uniquely identified in young and old supernatants, respectively. A total of 5 proteoglycans were identified in both young and old cohorts’ supernatants and a total of 11 matrisome associated proteins (25%). Interestingly, we identified lysyl oxidase (LOX) uniquely in young samples and hemopexin (HPX), antithrombin-III (SERPINC1), protein S100-A9 (S100A9) and transforming growth factor beta-2 (TGFB2) in old supernatants (**Table 3**).

**Table 3.**
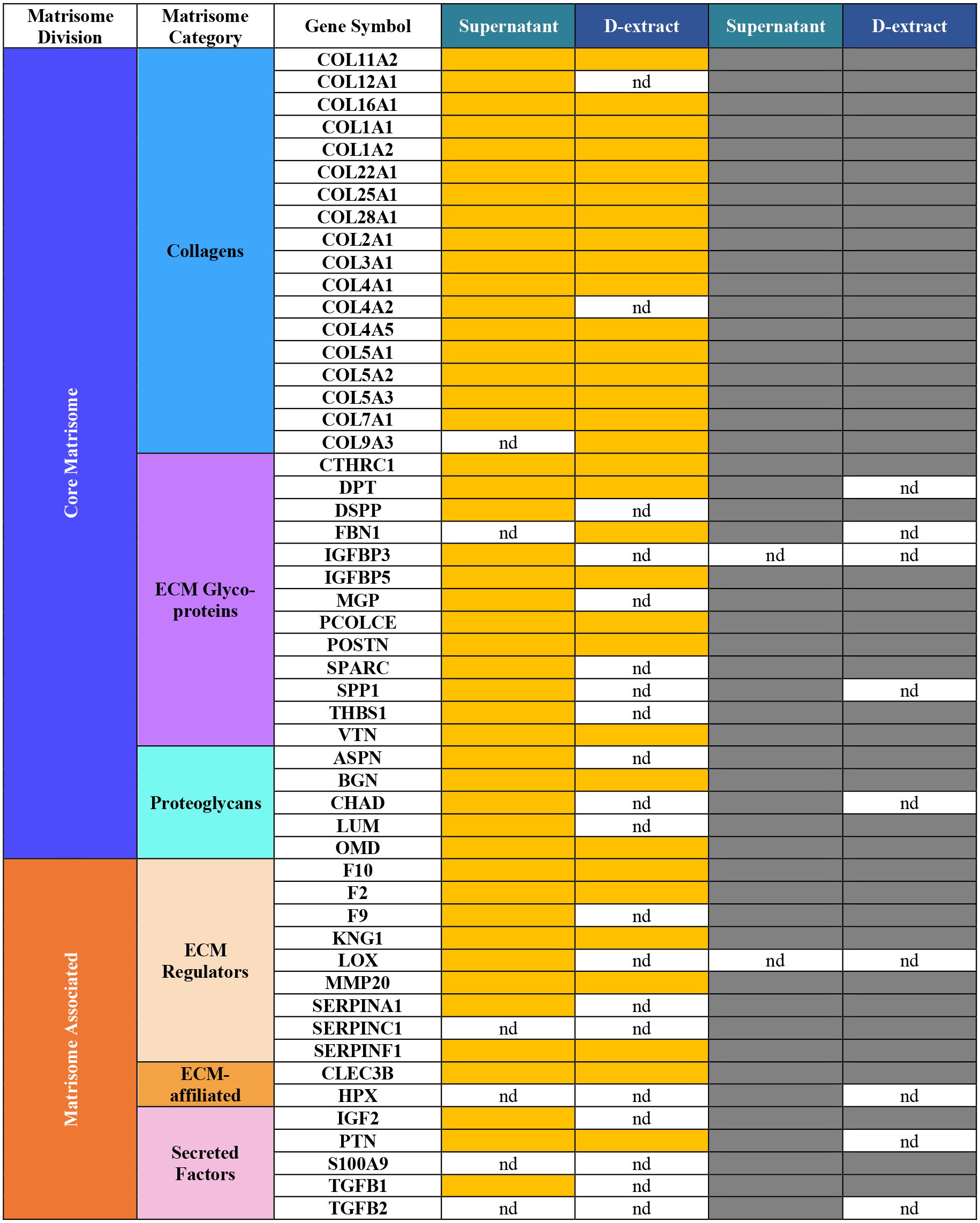
Comparing the young and old dentin matrisome. The proteins included in the list were detected with at least 2 peptides and in one or more individuals. Data are sorted by extracts and matrisome categories (nd: not detected).

Variability of the identified ECM proteins increased with age in the more insoluble fraction (D-extract), where 29 ECM proteins were found in both age cohorts, approximately 20% less common ECM proteins when compared to common proteins in the supernatants. A higher number of unique ECM proteins was identified in old D-extracts (14) when compared to young (3) (**Fig. 4C**). There was an increase in the number of identified ECM glycoproteins, proteoglycans and ECM regulators in D-extracts with age. Collagens such as COL12A1 and COL4A2, ECM glycoproteins like dentin sialophosphoprotein (DSPP), matrix gla protein (MGP), secreted protein acidic and cysteine rich (SPARC), and THBS1, and proteoglycans asporin (ASPN) and lumican (LUM) were found exclusively in old D-extracts, while fibrillin 1 (FBN1), osteopontin (SPP1), and dermatopontin (DPT) were uniquely observed in young D-extracts (**Table 3**).

#### 3.6.3. Proteomics provides potential insights into structural changes of dentin ECM proteins during aging

Bottom-up proteomics, which infers protein identity based on the identification of peptides, has also the potential to shed light on protein structure via peptide location fingerprinting (Eckersley et al., 2018, 2020). Indeed, structural changes of long-lived proteins, like ECM proteins, for example during tissue aging, can result in altered proteolytic susceptibility which can be assessed by evaluating protein-sequence coverage (**Supplementary Table 1C, columns AP to BG**) and mapping identifying peptides to primary protein structures. Here, we sought insight hinting at possible changes in ECM protein structures as a function of age. While our sample size limited our statistical analysis, we could identify ECM proteins whose coverage, and hence possibly structure, varied with age.

Biglycan (BGN) has previously been associated with mineralization (Haruyama et al., 2009). We detected peptides that cover on average 38% of the protein sequence in young insoluble dentin samples (D-extracts), while the sequence coverage only reached 14.5% on average across older insoluble dentin samples (**Fig. 5 and Supplementary Table 1C, row 35, and Supplementary Table 1N**). Conversely, we detected peptides covering on average 18.3% of the protein sequence in young EDTA-soluble dentin samples, while the sequence coverage only reached 27.2% on average across older EDTA-soluble dentin samples (**Supplementary Table 1C, row 35, and Supplementary Table 1N**). Moreover, we observed peptides that were either uniquely identified in young D-extracts or old D-extracts (**Fig. 5**). As the exclusive detection of these peptides in one sample type or the other could indicate greater accessibility to trypsin, we propose that a study on a larger number of samples may reveal structural differences that occur with the aging process.

**Figure 5.**
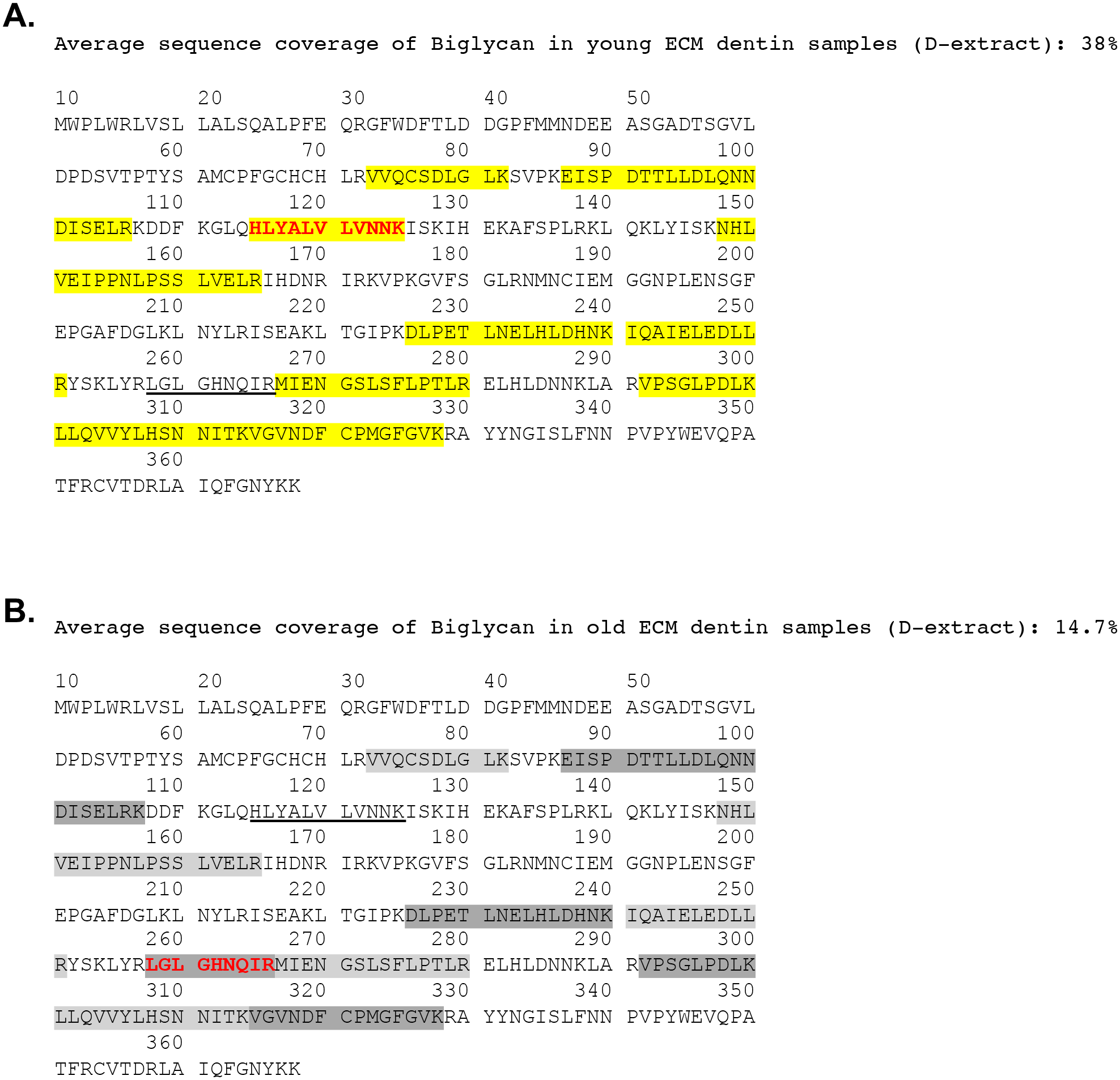
Sequence coverage map of biglycan in young and old dentin samples. Yellow and gray highlights on the biglycan sequence depict the peptides detected by mass spectrometry in young (**A**) and old (**B**) insoluble ECM dentin samples (D-extract). Red sequence highlights depict peptide differentially detected in young and old samples. Average sequence coverage of biglycan in young insoluble ECM dentin samples (D-extract) reaches 38%, while it averaged 14.7% in old insoluble ECM dentin samples *(**see*** ***Supplementary Table 1N***).

Another such example is the α2 chain of type V collagen (COL5A2). We detected peptides covering on average 12.8% of the protein sequence in young insoluble dentin samples, while the sequence coverage only reached 7.8% on average across older insoluble dentin samples. Conversely, we detected peptides covering on average 3.8% of the protein sequence in young EDTA-soluble dentin samples, while the sequence coverage only reached 8.4% on average across older EDTA-soluble dentin samples (**Supplementary Table 1C, row 9**).

Insulin-like growth factor binding protein-5 (IGFBP5) is known for its role in regulating cell growth, proliferation, and differentiation in different tissues, specifically in osteoblasts and odontoblasts (Aizawa et al., 2019; Mukherjee & Rotwein, 2008). We detected peptides covering on average 13.7% of the protein sequence in young EDTA-soluble dentin samples, while the sequence coverage only reached 4% on average across older EDTA-soluble dentin samples (**Supplementary Table 1C, row 26**).Results also reveal a relatively low coverage of minor ECM components of the dentin, as compared to large, structural, ECM proteins, such as the α1 and α2 chains of type I collagen whose sequence coverages reaches on average 70% across fractions. This highlights how much more remains to be there is to learn about the dentin matrisome.

## 4. Discussion

In this study, we demonstrate the feasibility of using proteomics to characterize the ECM composition of human cervical root dentin from single individuals and identify age-related differences in the composition and even possibly the structure of dentin ECM proteins.

In order to acquire an adequate amount of material, previous dental proteomic studies pooled samples from different samples (Abbey et al., 2018; Jágr et al., 2012, 2016, 2019; Park et al., 2009; Salmon et al., 2013). The protein analysis of small biological samples, such as tissue fragments and cells are an important challenge in the field of proteomics, however, advances in sample preparation and MS/MS techniques enabled highly effective studies and protein identification. Here, we show that the amount of material obtained from the cervical area of root dentin from one tooth, and thus from a single individual, is sufficient to conduct a rigorous proteomic analysis.

Prior studies have proposed that analyzing pools of ECM proteins of high, intermediate, and low solubility from tissue could shed light on their degree of assembly in the ECM scaffold (most insoluble fraction) and hence on their structural and functional properties (Barrett et al., 2018; Schiller et al., 2017). Indeed, ECM protein solubility is an important factor of ECM homeostasis and biology and can further influence the mechanical properties of tissues. Multiple mechanisms contribute to determining the level of assembly (or insolubility) of ECM proteins, including post-translational modifications such as cross-linking or, on the contrary, protein cleavage or degradation (Madzharova et al., 2019; Rogers & Overall, 2013). Here, we applied such approach, for the first time to the study of a mineralized tissue and found that protein fractions of different solubility have different biochemical composition.

Our study design also enabled the assessment of the extent of inter-individual variability within cohorts of the same age group. Our hypothesis was that variability within the ECM protein composition of young root dentin is limited. Nonetheless, it is important to acknowledge that the human oral microbiome is highly singular, and can be affected by lifestyle, hygiene, environment, genetics, diet, and disease (Willis & Gabaldón, 2020). Studies of mineralized tissues such as bone have shown that the effects of fluoride intake (Robinson & Kirkham, 1990), diet and the acid/base balance can lead to changes in tissue development, i.e., alterations in the abundance of collagen in the organic matrix (Balasse et al., 1999; Seibel, 2005). Therefore, a certain variability among the ECM protein composition of different individuals was anticipated. As an effort to limit this variability, samples were carefully selected to be within stringent age ranges and of same gender. Results show limited inter-individual variability and comparable protein composition between extracts of similar biochemical properties. Of note, studies on bone, another mineralized tissue, have found differences between samples of female and male individuals and indicate that inter-individual variability could be associated to hormonal differences between samples of different gender (Almeida et al., 2017). It would thus be interesting, in the future, to expand this study to samples from female individuals.

Comparison of the data from this study with previous studies revealed a substantial similarity between human dentin proteins. Yet, we report here the identification of a set of 13 ECM proteins never yet described in the human root dentin. The differences observed in the number of identified ECM proteins shown could be justified by methodological differences including sample collection and preparation, rigor of protein identification and annotation methods. The methodology employed here is tailored to specifically study ECM proteins, while other studies employed more generic proteomics pipeline. In addition, ECM proteins, and collagens in particular, present a specific set of post-translational modifications, namely hydroxylation of lysines and prolines that must be allowed as variable modifications, if, one wants to identify collagen peptides in mass spectrometry datasets (Naba et al., 2017; Taha & Naba, 2019). Of note, our starting material was the cervical region of the root dentin, whereas previous studies utilized the entire root (Jágr et al., 2012; Park et al., 2009). Structural and chemical differences have been identified in different regions of the root (Balooch et al., 2001; Koester et al., 2008), thus regional compositional variations is expected.

The comparison of the young and old human dentin root matrisome identified 2 proteins exclusively in young samples and 4 in old samples, all of which are of interest due to their potential roles in aging. The insulin-like growth factor binding protein 3 (IGFBP3) and lysyl oxidase (LOX) were uniquely identified in young samples. IGFBP3 is an insulin-like growth factor binding protein and recognized to function in the circulation (Baxter, 2000), with age, studies have shown that IGFBP3 appears to be decreased (Blaney Davidson et al., 2005; Hong & Kim, 2018). LOX is an enzyme that has an active role in initiating covalent cross-link formation in collagen fibrils (Kagan & Li, 2003). Its expression is associated to advanced glycation end products (AGEs), age-related modifications to bone mineralization and differentiation of cells in dental tissues (Aoki et al., 2013; Kim et al., 2012). The 4 proteins uniquely identified in old samples are hemopexin (HPX), SERPINC1, S100A9 and transforming growth factor beta-2 (TGFB2). S100A9 is a calcium-binding protein (Zreiqat et al., 2007) implicated in receptors for AGEs (Hiroshima et al., 2018; Xu et al., 2013), alveolar bone resorption and a mediator of aging-associated disorders (Maekawa et al., 2019; Swindell et al., 2013). TGFβ2 is a ligand of the transforming growth factor-beta superfamily (Baxter, 2000) and disruptions of the TGFβ/SMAD pathway has also been linked to disease and age-associated disorders (Tominaga & Suzuki, 2019). HPX and SERPINC1, proteins linked to the circulation and recently associated to Alzheimer’s disease (Ashraf et al., 2020) and aging of the skin (Ma et al., 2020), but whose roles in the dentin remain to be understood.

Previous metabolic and proteomic investigations in different model organisms such as mice (Angelidis et al., 2019), nematodes (Copes et al., 2015), and humans (Chaleckis et al., 2016), have shown alterations in ECM abundance with age, however, the underlying mechanisms are yet to be determined. In addition, shifts in protein solubility with age have been shown in protein profiles of young and old superficial digital flexor tendons (Peffers et al., 2014) and between lung samples of young and old mice (Angelidis et al., 2019). However, the functional implications and cause of these changes are still unknown. We suggest that changes in the composition of dentin ECM with age, regarding proteins that are linked to mineralization, collagen post-translational modifications and the accumulation of AGEs could potentially play important roles in alteration of the dentin ECM ultrastructure, mineral and organic content, biomechanical performance and consequently the functionality of teeth with age. Further proteomic investigations using label-based quantitative methods, or focusing on post-translational modifications, and the functional roles of specific proteins may reveal additional differences occurring in the dentin ECM during aging.

## 5. Conclusions

This study provides a robust experimental framework to explore the composition of the root dentin ECM. Proteomic analysis from single dental roots revealed a feasible profiling of the dentin ECM with limited inter-individual variability among young samples. The analysis of the matrisomes of young and old dentin root samples showed age-related changes to the solubility and composition of the dentin ECM. Future investigations are now necessary to unveil age-dependent mechanisms leading to these changes. In addition, beyond protein identification, we demonstrate the potential of proteomics into gaining structural insight on dentin ECM proteins. With the development of refined methods capable of revealing quantitative (*e.g.,* TMT-labeling) and structural (*e.g.,* degradomics, peptide and PTM mapping on 3D protein structures) information, this proteomic pipeline can become a method of choice for future studies focused on the matrisome of the tooth-periodontal-bone complex.

## CRediT author statement

**Mariana Reis:** Methodology, Validation, Formal analysis, Investigation, Writing - Review & Editing.

**Fred Lee:** Methodology, Validation, Formal analysis, Investigation, Writing - Review & Editing.

**Ana K. Bedran-Russo:** Conceptualization, Methodology, Formal analysis, Investigation, Resources, Writing - Review & Editing, Visualization, Supervision, Project administration, Funding acquisition.

**Alexandra Naba:** Conceptualization, Methodology, Formal analysis, Investigation, Resources, Writing - Review & Editing, Visualization, Supervision, Project administration, Funding acquisition.

## Acknowledgements

The authors would like to thank Dr. Hui Chen from the Mass Spectrometry Core facility at the University of Illinois at Chicago and the Dr. George Chlipala from the Research Informatics Core facility at the University of Illinois at Chicago for their technical assistance.

## Supplementary data

**Supplementary Table 1. Complete mass spectrometry dataset**

**Supplementary Table 2. Comparison of the matrisome of young dentin samples from multiple studies**

**Supplementary Figure 1. Comparison of the matrisome of young dentin samples from multiple studies**

---

**Figure S1.**
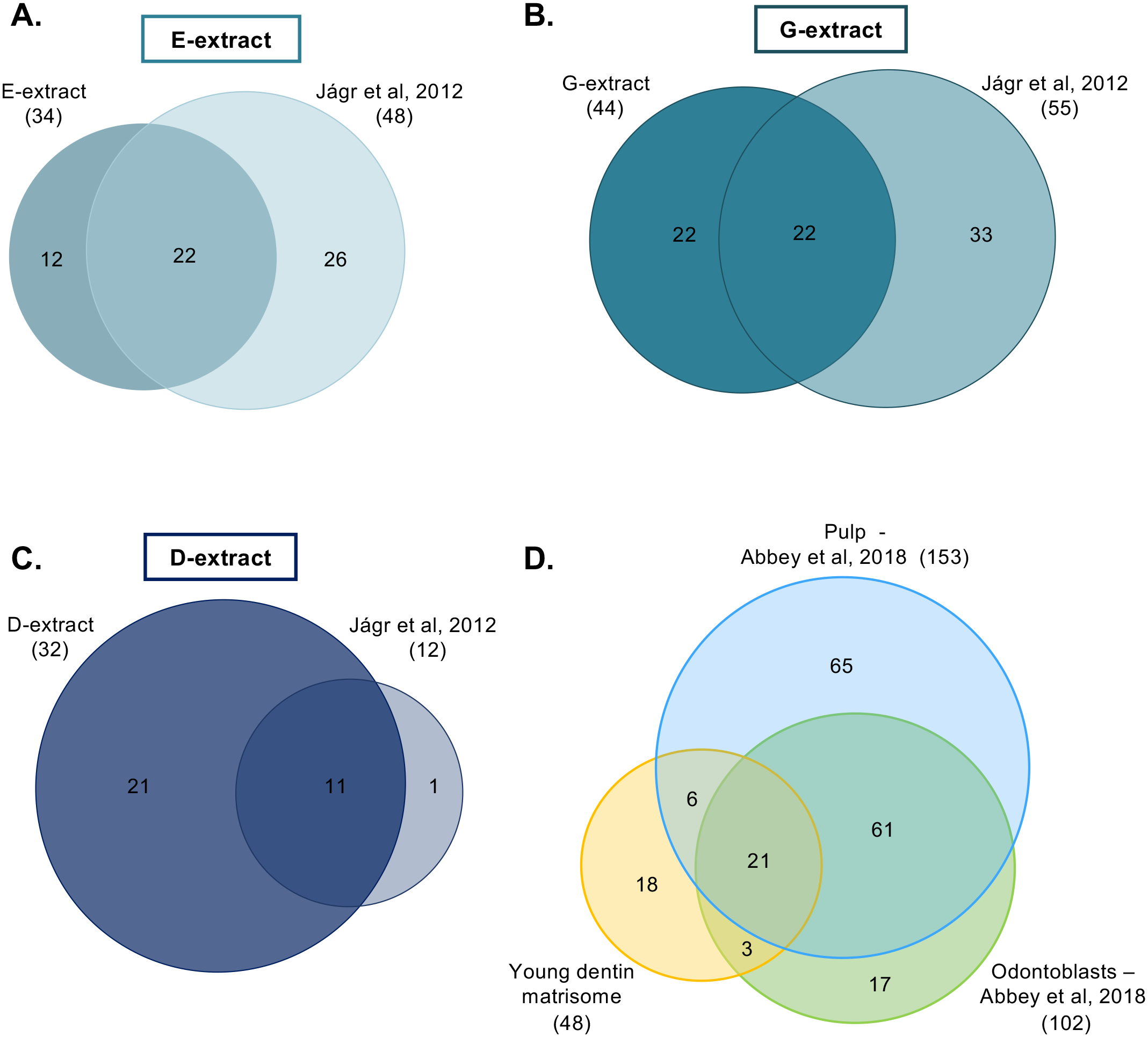
Comparison of the matrisome of young dentin samples from multiple studies. Comparison of the composition of E-extracts **(A)**, G-extracts **(B)**, and D-extracts **(C)**. The complete matrisome of each study is presented in Supporting Table 3, where the proteins are grouped into matrisome divisions and categories. Venn diagrams with the comparison of proteins identified in **(D)** the young dentin matrisome, pulp and odontoblast layer from Abbey et al., 2018. *Related to Supplementary Table 2*.

